# Tracing developmental and adult hematopoiesis with an endogenous zebrafish *runx1-2A- CreERT2* CRISPR knock-in

**DOI:** 10.64898/2026.07.03.736368

**Authors:** James A. Preston, Masuma K. Usha, Stephen C. Ekker, Karl J. Clark, Jeffrey J. Essner, Raquel Espin-Palazon, Maura McGrail

## Abstract

Zebrafish combines the power of genetics and unparalleled *in vivo* imaging to investigate the dynamics of vertebrate hematopoietic development. The conserved transcription factor Runx1 is essential for definitive hematopoiesis. We generated an endogenous zebrafish *runx1-2A-creERT2* CRISPR knock-in for tamoxifen-regulated Runx1 lineage tracing and characterized its activity using recombinase-dependent fluorescence reporters, live imaging and flow cytometry. Tamoxifen induction of Cre in the early embryo labeled all expected Runx1 lineages, including neuroectodermal olfactory placode and Rohan-Beard neurons, primitive hematopoietic blood cells, and emerging definitive hematopoietic stem and progenitor cells (HSPCs) in the dorsal aorta. Short windows of tamoxifen treatment allowed separation of Runx1 primitive from definitive hematopoiesis and visualization of HSPC colonization of the larval caudal hematopoietic tissue, thymus and kidney. Runx1 HSPCs labeled during development persisted into adulthood and contributed to hematopoietic precursors, myeloid, lymphoid, and peripheral blood lineages. Labeling of all blood lineages was also effective by tamoxifen treatment of 5-month-old adults. The *runx1-2A-creERT2* knock-in line provides a novel genetic tool for inducible spatial and temporal analysis of Runx1 progenitor mechanisms in developmental and adult hematopoiesis.

**Summary Statement:** Zebrafish *runx1-2A-creERT2* knock-in provides inducible spatial and temporal Cre recombinase Runx1 lineage labeling in development and adults, and reveals embryonic Runx1 precursors contribute to adult hematopoiesis.

## Introduction

The accessibility of the zebrafish embryo to *in vivo* imaging with HSPC fluorescent reporter lines has provided remarkable insight into the developmental dynamics of vertebrate hematopoiesis ^1–7^.

However, hematopoietic stem cell-specific tools for inducible, spatial and temporal HSPC imaging and functional genetic analysis in zebrafish are unavailable. Existing reporter and driver transgenics are either broadly expressed in mesoderm-derived endothelial cells of the vascular or lymphatic blood vessels, or contain incomplete gene regulatory elements inherent in cloned promoter/enhancer constructs. To overcome these limitations in traditional zebrafish transgenesis, we recently established CRISPR precision targeted integration ^8,9^ in zebrafish and applied this method to generate Cre recombinase knock-ins in lineage defining transcription factors that recapitulate the endogenous, tissue-specific expression profile of the gene ^10^. We demonstrated endogenous Cre and tamoxifen inducible CreERT2 drivers enable lineage-specific *in vivo* progenitor tracing ^10,11^ and cell autonomous Cre recombinase-mediated genetic analysis of progenitor proliferation ^12^. Here, we describe an endogenous CreERT2 CRISPR knock-in in the definitive hematopoietic marker Runx1 and demonstrate its activity in spatial and temporal tracing of Runx1 lineages in developmental and adult hematopoiesis.

Blood development in the vertebrate embryo is driven by highly conserved mechanisms that specify vascular endothelial progenitors in the gastrula lateral plate mesoderm ^13–17^. Primitive hematopoiesis in the mammalian yolk sac ^18^ and teleost intermediate cell mass ^19^ gives rise to early embryo erythromyeloid cells. Definitive HSPCs arise from aortic endothelial cells of the mammalian aortic- gonadic-mesonephros (AGM) ^20^ and the zebrafish AGM equivalent the dorsal aorta (DA) ^21^ through endothelial-to-hematopoietic transition (EHT) ^22–24^. In mammals, HSPCs colonize the fetal liver and thymus and seed the adult hematopoietic niche in the bone marrow ^18,25^, whereas in zebrafish HSPCs migrate to the larval caudal hematopoietic tissue (CHT) ^24,26^ followed by colonization of the thymus and the adult hematopoietic kidney ^3,24,27^. The long-standing view in the field proposes that HSPCs born in the embryo maintain self-renewing potential throughout life to generate adult blood ^20^.

However, recent work in mice shows embryonic multilineage progenitors also persist into adulthood and contribute to blood production ^28^.

The definitive hematopoietic transcription factor Runx1 ^22,29–31^ is essential for HSPC emergence from hemogenic endothelium ^31–33^. In the zebrafish gastrula *runx1* is first expressed in the lateral plate mesoderm ^30,34,35^ and the anterior neural plate ^34^. Later, *runx1* is found in both mesodermal and neural derivatives ^30,35^. Multiple zebrafish *runx1* promoter:fluorescent reporter transgenics have faithfully mapped *runx1* lineages to the neural olfactory placode and Rohan Beard cells, and hematopoietic cells in the DA, CHT, thymus, pronephros ^2–4,6^ and adult kidney marrow ^4,36^. In adult mice *Runx1* is broadly expressed in HSPCs, myeloid cells and a subset of lymphoid lineages ^37,38^. Conditional *Runx1* knock-out in the adult mouse hematopoietic compartment resulted in a mild myeloproliferative phenotype and defect in platelet production ^39^, while lineage-specific knockout demonstrated a role in generating common lymphoid progenitors and B- and T-cell maturation ^39^. These studies underscore the critical role of Runx1 in definitive hematopoiesis during development and in adult hematopoietic lineage differentiation. However, currently available animal models do not enable temporal and spatial analysis of Runx1 hematopoietic lineages spanning the embryo-to-adult transition.

To generate a novel zebrafish line for inducible Runx1 lineage tracing and functional gene studies, we isolated an in frame *runx1* CRISPR knock-in that drives expression of tamoxifen-regulated Cre recombinase CreERT2. Using live confocal imaging we demonstrated *runx1* CreERT2 labeling with a floxed reporter recapitulated all Runx1 progenitor-derived neuroectodermal and mesodermal hematopoietic lineages in the early embryo. Runx1 progenitors contributed to primitive hematopoietic erythromyeloid cells and definitive HSPCs. Time lapse imaging showed Runx1-labeled cells budded from the DA hemogenic endothelium and moved into circulation through the posterior caudal vein.

Runx1 HSPCs populated the caudal hematopoietic tissue and later colonized the larval thymus and pronephros region. Tamoxifen induction of *runx1* CreERT2 lineage tracing at distinct time points during development distinguished hematopoietic populations arising from primitive and definitive hematopoiesis and colonization of Runx1 HSPCs to thymus and kidney. Long term and adult lineage tracing demonstrated embryo Runx1 progenitors contribute to adult hematopoiesis, giving rise to all blood lineages. The *runx1-2A-creERT2* line provides a powerful genetic tool to further investigate *runx1*-expressing lineages and cell autonomous mechanisms of cell fate during embryonic and adult hematopoiesis.

## Design

Hematopoietic stem cell-specific tools for inducible, spatial and temporal HSPC imaging and functional genetic analysis in zebrafish are currently unavailable. Existing zebrafish HSPC reporter and driver transgenics in use are broadly expressed in endothelial cells of the vascular or lymphatic blood vessels, and generated with Tol2 or BAC transgenesis of promoter:cDNA constructs. Here, we describe a novel zebrafish endogenous *runx1* CreERT2 CRISPR knock-in isolated with our highly efficient GeneWeld precision targeted integration technology ^8,9^. Our GeneWeld approach uses a plasmid based template and CRISPR-Cas9 short homology directed targeted integration to direct knock-in of a *2A-creERT2* cassette linked to a tissue specific secondary marker for easy allele identification ^10,11^. We demonstrate *runx1-2a-creERT2* activity enables Runx1-specific lineage tracing of embryonic progenitors and HSPCs in developmental and adult hematopoiesis.

## Results

### A Zebrafish *runx1-2A-creERT2* knock-in line generated with GeneWeld CRISPR targeted integration

To recover a zebrafish line expressing CreERT2 under the control of *runx1* endogenous gene regulatory elements, GeneWeld CRISPR/Cas9 homology directed targeted integration ^8,9^ was used to knock-in a *2A-creERT2* cassette at the 3’ end of the coding sequence in *runx1* (ZDB-GENE-000605- 1) (Figure 1A, B; Figure S1A; Table S1) following our previously published method ^10,11^. The GeneWeld *2A-creERT2*, *exorh:CFP* knock-in cassette contained a linked secondary marker with the *exorh* promoter ^40^ driving pineal-specific Cerulean Fluorescent Protein (CFP) expression, allowing allele identification by fluorescence screening on a stereomicroscope (Figure 1C). The cassette was integrated 29 bp upstream of the *runx1* exon 7 translation stop codon (Figure 1A, Figure S1B). The targeting vector contained a 5‘ homology arm with mutated PAM and the remainder of the *runx1* coding sequence (Materials and Methods). One precise integration line was used to establish the *runx1-2A-creERT2, exorh:CFP^is97^* line (Figure S1B), referred to as *runx1-2A-creERT2*. Development and viability appear normal in *runx1-2A-creERT2* heterozygotes and homozygotes.

**Figure 1.**
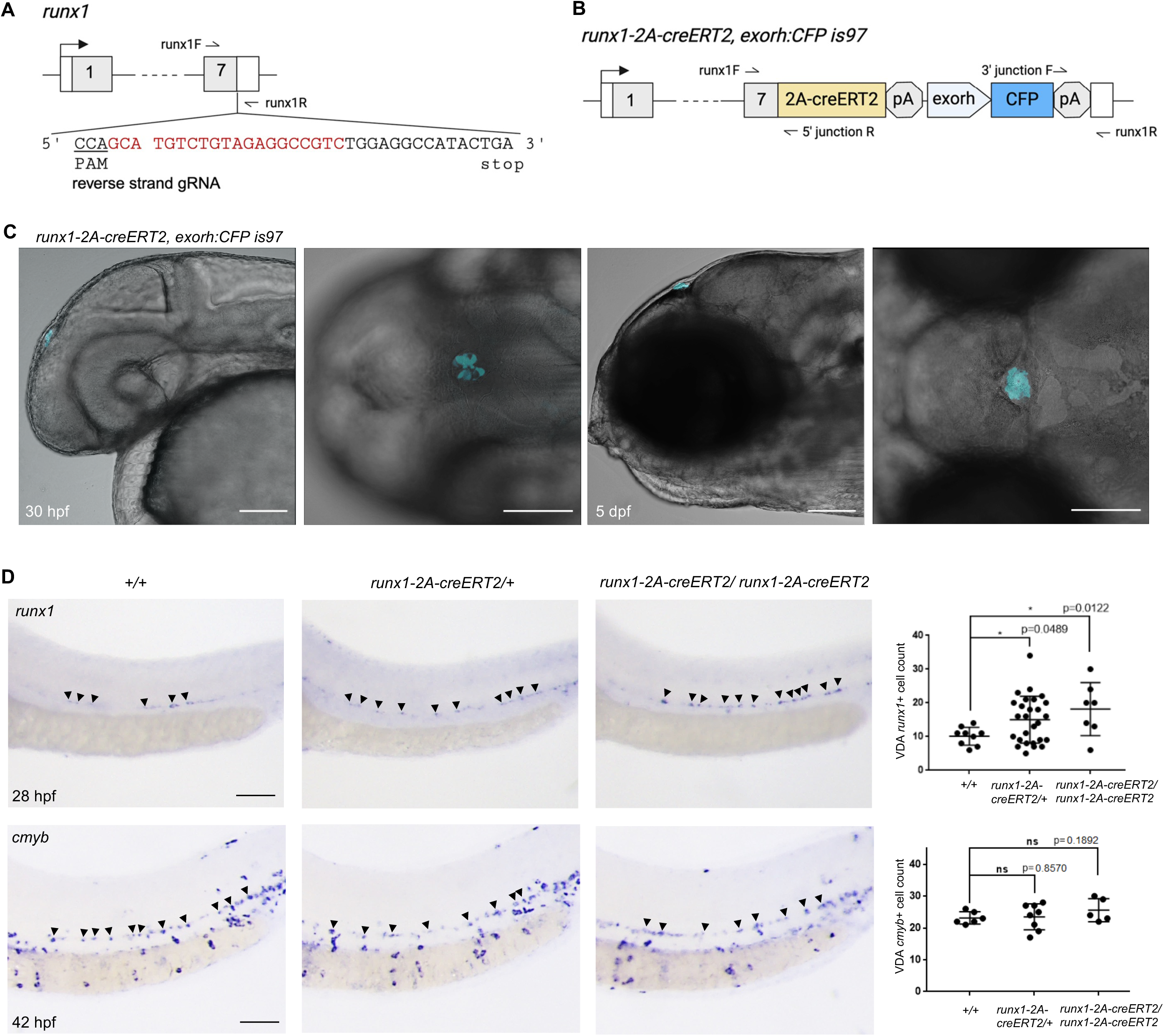
The zebrafish *runx1-2A-creERT2* knock-in line and analysis of hematopoiesis in heterozygous and homozygous embryos. **A** Zebrafish *runx1* gene diagram with location of exon 7 CRISPR/Cas9 gRNA and targeted integration site. **B** Diagram of *runx1-2A-creERT2, exorh:CFP^is97^* Knock-in line with cassette integrated in frame at the 3’ end of the *runx1* coding sequence. Additional information in Figure S1. **C** Lateral and dorsal views of *runx1-2A-creERT2/+* 30 hpf embryo and 5 dpf larva confocal live imaging showing pineal-specific CFP expressed from the *exorh:CFP* transgenesis marker. **D** Lateral trunk views of 28 hpf *runx1*-hybridized and 42 hpf *cmyb*-hybridized +/+, *runx1-2A- creERT2*/+ and *runx1-2A-creERT2/runx1-2A-creERT2* embryos. Arrowheads point to *runx1+* and *cmyb+* hemogenic endothelial cells in the DA. Anterior is to the left, Dorsal is up. Plots show quantification with SEM of *runx1+* cells in +/+ (n=9), *runx1-2A-creERT2*/+ (n=27), and *runx1-2A- creERT2/runx1-2A-*creERT2 (n=7) embryos, and *cmyb+* cells in +/+ (n=6), *runx1-2A-creERT2*/+ (n=8), and *runx1-2A-creERT2/runx1-2A-*creERT2 (n=5) embryos. Scale bars 100 μm.

### HSPC Specification is maintained in *runx1-2A-creERT2* embryos

To determine whether integration of the *2A-creERT2, exorh:CFP* cassette disrupted *runx1* expression or HSPC specification in the Dorsal Aorta (DA), *in situ* hybridization was performed on 28 hpf and 42 hpf *runx1-2A-creERT2* embryos with probes for *runx1* and the HSC proliferation regulator *cmyb* ^41^ (Figure 1D). The number of DA positive cells was quantified in heterozygous *runx1-2A-creERT2/+,* homozygous *runx1-2A-creERT2/runx1-2A-creERT2*, and wild type control sibling embryos. At 28 hpf, the number of DA *runx1*+ cells in heterozygotes and homozygotes was higher than in wild type, which may result from an increase in mRNA stability due to the strong ocean pout transcriptional terminator ^10,42^ following the *creERT2* cDNA. However, DGE analysis of bulk RNASeq from 28 hpf wildtype embryos compared with either heterozygous or homozygous *runx1-2A-creERT2* showed no significant difference in expression across Runx1 target genes and downstream pathways regulating EHT and definitive hematopoiesis (Figure S1C; Table S2). At 42 hpf the number of DA *cmyb*+ HSPCs was not significantly different in either *runx1-2A-creERT2* heterozygotes or homozygotes. Together, these data indicate the *runx1-2A-creERT2* integration did not negatively impact specification of DA hemogenic endothelium HSPCs.

### Inducible Runx1 progenitor lineage tracing in zebrafish embryo and larva

Tamoxifen inducible recombinase lineage tracing was performed on *runx1-2A-creERT2* embryos carrying the ubiquitous floxed switch reporter *ubb:lox-eGFP-Stop-lox-mCherry* (*ubi:Switch*) ^43^. Cre recombination excises the GFP-Stop sequences and leads to a permanent switch from GFP to mCherry expression. *runx1-2A-creERT2; ubi:Switch* embryos were treated with 5 μM 4-OHT continuously from shield stage until imaging at 24 hpf, 48 hpf, 72 hpf and 5 dpf (Figure 2A). *runx1-2A- creERT2; ubi:Switch* shield stage embryos treated with EtOH vehicle alone did not show detectable mCherry expression at 28 hpf or 48 hpf (Figure S2A, B).

**Figure 2.**
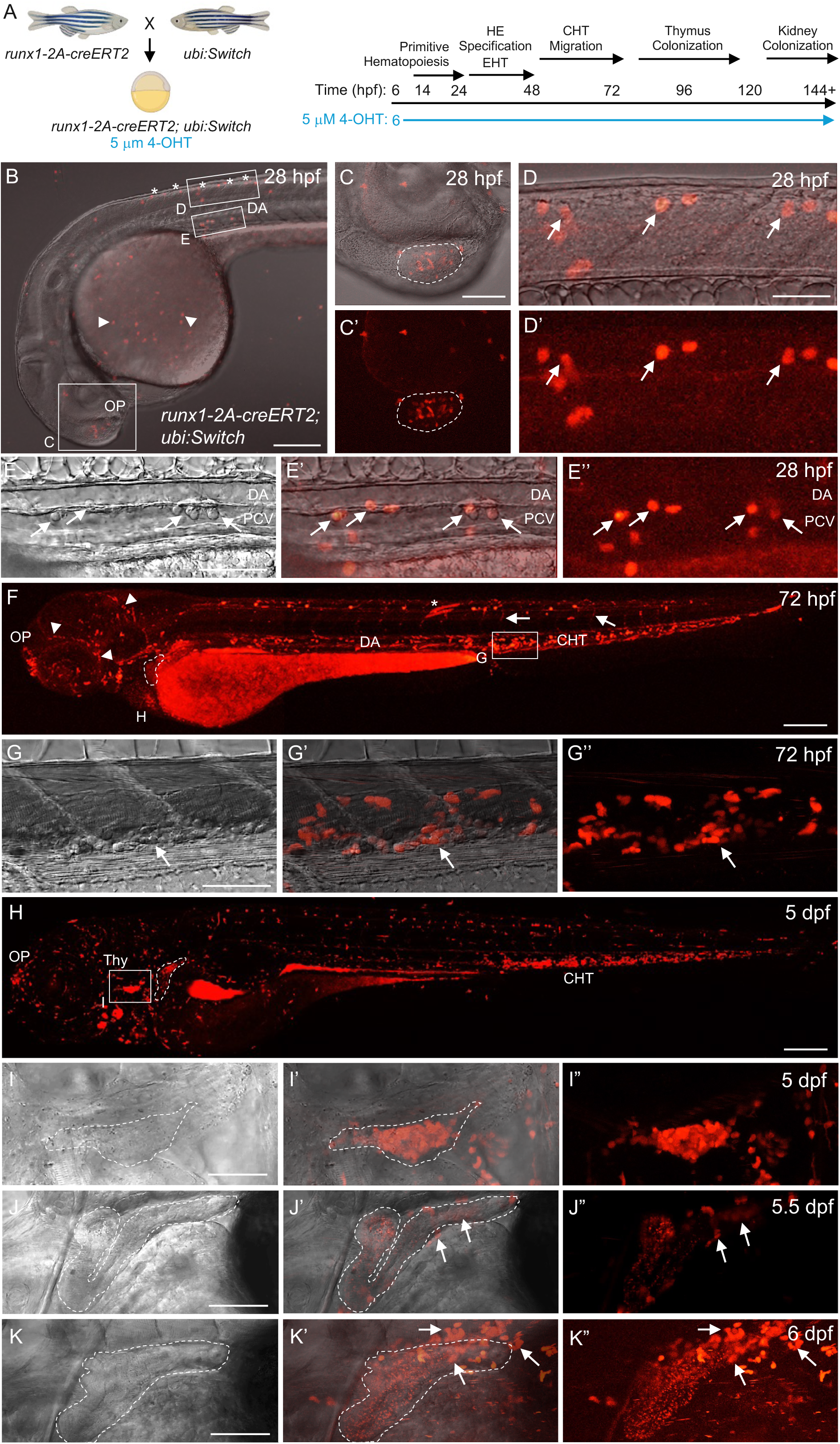
Tamoxifen induced *runx1* lineage tracing in *runx1-2A-creERT2; ubi:Switch* embryos and larva after shield stage 4-OHT treatment. **A** Schematic showing genetic cross to generate *runx1-2A-creERT2; ubi:Switch* embryos for shield stage 4-OHT induction and timeline of critical events in hematopoiesis. **B – I** Live confocal imaging of mCherry expression in *runx1* lineages in embryos and larva. **B** 28 hpf embryo with mCherry in OP, Rohon-Beard neurons (asterisks), macrophages (arrowhead), and DA. Panel B red channel brightness digitally increased 40% to reveal mCherry^+^ pattern. **C, C’** Higher magnification view mCherry^+^ cells in the OP (dashed line). **D, D’** Higher magnification lateral view of mCherry^+^ Rohon-Bearn Neurons (arrows) in the spinal cord. **E - E’’** Higher magnification lateral view of mCherry^+^ cells (arrows) budding from the ventral DA. **F** 72 hpf larva with mCherry^+^ cells in OP, cranial blood vessels (arrowheads), heart, DA, CHT, trunk skeletal muscles (asterisk), and intersegmental vessel (arrows). Background labeling present in proximal convoluted tubule (dashed line) of the anterior pronephros. **G - G’’** Higher magnification view of mCherry^+^ cells in the CHT (arrow) at 72 hpf. **H** 5 dpf larva with mCherry^+^ cells in OP, Thy, and CHT. Anterior pronephros outlined (dashed line). **I – I”** Higher magnification images of mCherry^+^ cells in the developing thymus (dashed outline). **J – J”** mCherry^+^ cells (arrows) in the anterior pronephros tubules (dashed line) and adjacent tissue at 5.5 dpf. **K – K”** mCherry^+^ cells (arrows) in tissue surrounding the anterior pronephros tubules (dashed line) at 6 dpf. CHT, caudal hematopoietic tissue; DA, dorsal aorta; EHT, endothelial-to-hematopoietic transition; H, heart; HE, hemogenic endothelium; OP, olfactory placode; PCV, posterior caudal vein; Thy, thymus. n=3 embryos or larva imaged at each time point. Scale bars: B, F, H, 200 μm; C, D, E, G, I, J, K 50 μm.

*runx1-creERT2* lineage tracing recapitulated all expected Runx1 progenitor descendants in 28 hpf embryos ^30,35^ (Figure 2B-I). mCherry expression was detected in the neuroectoderm derived olfactory placode (Figure 2B, C) and neural plate derived Rohan-Beard neurons in the dorsal neural tube (Figure 2B, asterisks; 2D, D’ arrows). mCherry labeled cells were present throughout the head and around the yolk sac, which may represent primitive macrophages (Figure 2B, arrowhead), consistent with the requirement for murine *Runx1* in primitive erythromyeloid cells ^30,44^. In the dorsal aorta, mCherry^+^ cells were detected budding from the ventral wall into the subaortic space (Figure 2B DA; Figure 2E - E’’ arrows). Time lapse imaging shows mCherry^+^ cells bud from the DA and enter circulation after crossing into the posterior caudal vein (PCV) (Video S1, S2). Together, these results show the *runx1* lineage can be traced to neuroectodermal, primitive and definitive hematopoietic cells in the embryo, and is consistent with the requirement for *runx1* in HSPC emergence by EHT ^3,22^.

From 48 hpf to 72 hpf *runx1*-lineage labeling was maintained in olfactory placode cells and Rohan- Beard neurons (Figure 2F, Figure S2C). At 72 hpf an increase in mCherry was detected in the CHT (Figure 2F CHT, 2G-G”), and in the heart and cranial blood vessels (Figure 2F H, arrowheads), presumably representing circulating blood cells. Sparse labeling of trunk skeletal muscles (Figure 2F asterisk), and intersegmental vessels (Figure 2F arrow) as previously reported for the *runx1P2*:Citrine BAC transgenic line ^2^ was observed, and may result from multi-lineage progenitors that originate at the border between the Lateral and Intermediate Plate Mesoderm in the gastrula.

### Runx1 lineage tracing in larval thymus and kidney hematopoietic organs

In the 5 dpf larva *runx1* labeled cells were present in the thymus (Figure 2H, Thy; 2I – I”) and in the CHT (Figure 2H CHT). Time lapse imaging of the thymus in 4 dpf and 5 dpf larva showed mCherry cells within and migrating into and around the organ (Video S3, S4). Thymic labeling is consistent with previous reports of *runx1* promoter:fluorescent fusion protein transgenics labeling lymphoid precursors in the thymus from 3 – 5 dpf ^2–4^. In the developing pronephros, mCherry+ cells were first detected at 5.5 days in the tissue surrounding the anterior proximal tubules (Figure 2J-J”), occupying the perivascular region where previously described *runx+23:GFP* labeled anterior mediolateral HSPC clusters reside ^6^. Background punctate signal in the anterior proximal tubules at 72 hpf and 5 dpf (Figure 2F, H, J dashed outline) was variable across larva and also present in wildtype. The number of *runx1* mCherry+ cells populating the anterior pronephros increased at 6 days (Figure 2K-K”).

Together, these results demonstrate *runx1-2A-creERT2* enables effective lineage tracing of previously defined neuroectodermal and primitive and definitive hematopoietic Runx1 populations in the embryo. *runx1-2A-creERT2* labeling mapped hematopoietic precursors to the larval thymus and pronephros, which give rise to adult hematopoietic organs ^2,3,30,34^. Significantly, the CreERT2 cassette does not show ectopic activity in the absence of 4-OHT, and its activity faithfully recapitulates endogenous *runx1* expression, validating the line as a robust tool for temporally controlled lineage tracing of Runx1-expressing cells during development.

### Timed Cre recombinase induction distinguishes primitive and definitive Runx1 hematopoiesis

To distinguish *runx1*-lineage blood cells arising from primitive and definitive hematopoiesis, recombination switch labeling was induced at four hematopoietic transitions: from shield stage through primitive hematopoiesis, after onset of definitive hematopoiesis, early in larval CHT HSC expansion, and during migration of lymphoid precursors to the thymus. *runx1creERT2; ubi:Switch* embryos and larvae were treated with 5 μM 4-OHT for 24-hour intervals beginning at shield stage, 26hpf, 50hpf, and 74hpf (Figure 3A). Treatment of *runx1creERT2; ubi:Switch* from shield stage to 24 hpf (Figure S2D, E) recapitulated lineage labeling in embryos and larva after continuous treatment as shown in Figure 2. mCherry^+^ cells were present throughout the body, in blood vessels, and in the OP, RB, DA, and CHT at 48 hpf and 72 hpf (Figure S2D, E). Treatment from 25 – 48 hpf reduced the levels of mCherry^+^ RB cells in the spinal cord, and fewer mCherry cells were detected throughout the body (Figure S2F), consistent with the transition from primitive erythromyeloid hematopoiesis to definitive hematopoiesis. After 50-72 hpf treatment and imaging at 72 hpf (Figure S2G), or 74-96 hpf treatment and imaging at 5 dpf (Figure S2H), mCherry^+^ cells were predominantly present in the CHT and thymus.

**Figure 3.**
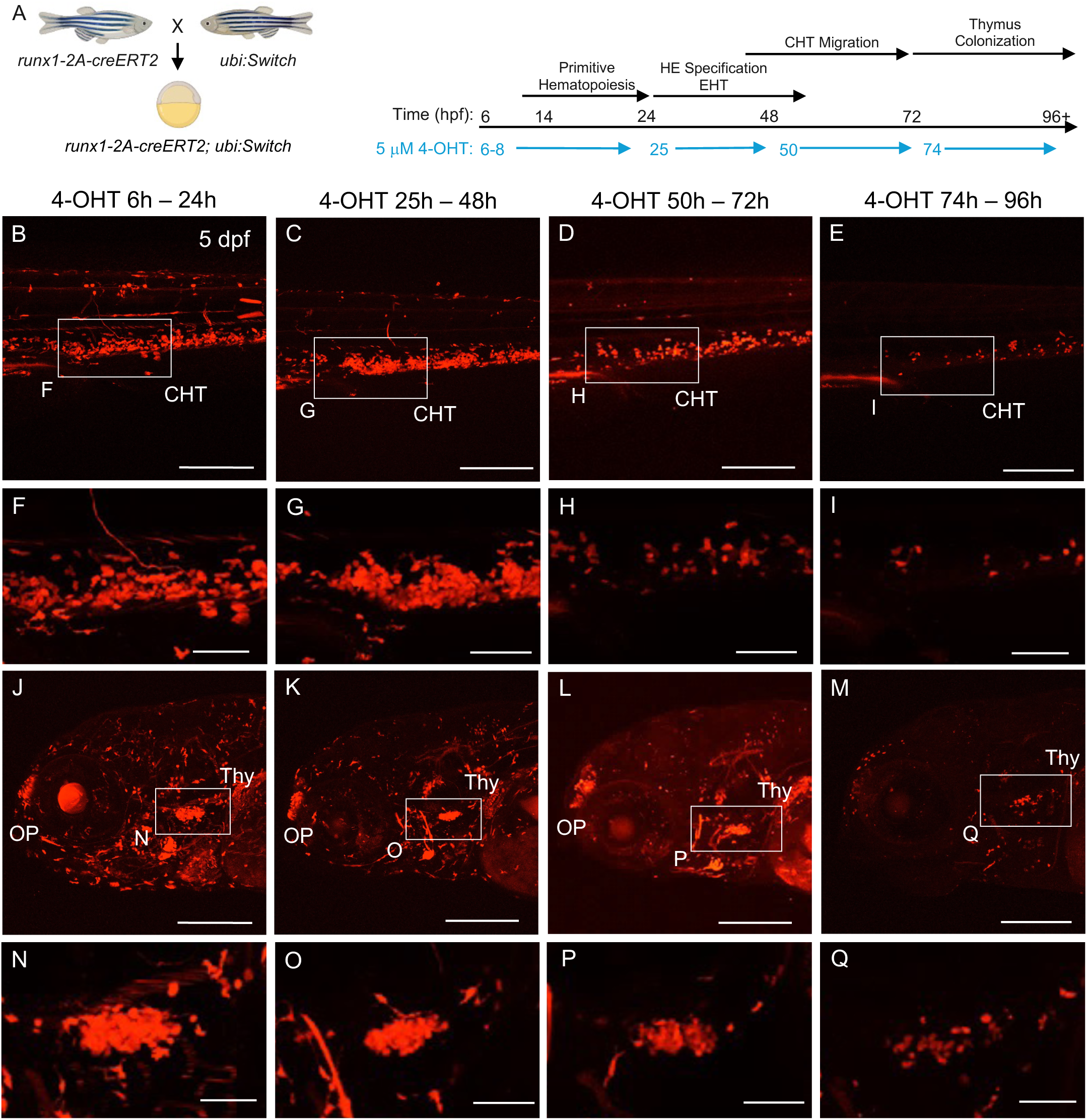
*runx1* lineage tracing in *runx1-2A-creERT2; ubi:Switch* larval thymus after 24 hour 4- OHT treatment windows beginning at shield stage, 1 dpf, 2 dpf, and 3 dpf. A Schematic showing genetic cross to generate *runx1-2A-creERT2; ubi:Switch* embryos and timeline of 4-OHT treatments during critical events in hematopoiesis. **B – I** Live confocal imaging of mCherry expression in 5 dpf *runx1-2A-creERT2; ubi:Switch* CHT after 4-OHT treatment from 6-24 hpf (**B**, **F**), 25-48 hpf (**C**, **G**), 50- 72 hpf (**D**, **H**), and 74-96 hpf (**E**, **I**). **J – Q** Live confocal imaging of mCherry expression in 5 dpf *runx1- 2A-creERT2; ubi:Switch* OP and Thy after 4-OHT treatment from 6-24 hpf (**J**, **N**), 25-48 hpf (**K**, **O**), 50- 72 hpf (**L**, **P**), and 74-96 hpf (**M**, **Q**). n=3 larva imaged at each time point. CHT, caudal hematopoietic tissue; EHT, endothelial-to-hematopoietic transition; HE, hemogenic endothelium; OP, olfactory placode; Thy, thymus. mCherry image brightness digitally increased in Powerpoint by 40% to reveal mCherry+ pattern for panels **B-E** and **J-M**. Scale bars B-E, J-M 200 μm; F-I, N-Q 50 μm.

Comparison of mCherry^+^ CHT cells in *runx1creERT2; ubi:Switch* 5 dpf larva suggested an initial increase after the 25-48 hpf switch that was reduced after later stage switching during day 2 and day 3 (Figure 3B-E, higher magnification images Figure 3F-I). The initial increase during day two correlates with the peak of HSPC specification in the DA ^24^. In the 5 dpf thymus, the relative level of mCherry^+^ signal decreased after switching during larval time points (Figure 3J-Q). The reduction in intensity agrees with previous reports of thymus seeding by HSC-independent lymphoid progenitors that originate in the aortic endothelium at 24 hpf ^45^ and later born CD41-GFP^+^ thymocytes ^46^. These results demonstrate robust 4-OHT temporal induction of *runx1-2A-creERT2* Cre activity can distinguish primitive from definitive hematopoiesis in the embryo and map colonization of definitive HSPCs to the larval CHT and thymus.

To examine colonization of the developing kidney by Runx1+ HSPCs derived from definitive hematopoiesis, *runx1-2A-creERT2; fli1:EGFP* adults were crossed with the *actb2:loxP-BFP-loxP- dsRed* ^23^ switch reporter to orient *runx1* dsRed+ cells relative to GFP-labeled vasculature surrounding the anterior pronephros (Figure 4A). Treatment of embryos and larva with 4-OHT in 24 hour intervals was performed beginning at shield stage, 25hpf, 50hpf, and 74hpf (Figure 4; Figure S3). Imaging of Runx1 dsRed+ blood cells recapitulated the pattern of CHT and thymus localization (Figure S3 A – O) described above for lineage tracing using the *ubi:Switch* reporter (Figure 2, 3). Induction of Cre activity during HSPC emergence from the DA resulted in dsRed+ cells in the region of the anterior pronephros (Figure 4 B – I) beginning at 5 dpf and increasing through 7 dpf. The dsRed+ cells were located in perivascular tissue surrounding the glomerulus, flanked by the posterior cardinal vein (PCV) and lateral dorsal aorta (LDA) (Figure 4 B – I), and in dorsal view appeared as two medial clusters of cells (Figure 4 F – I, dashed ovals). The timing, location and positioning relative to the vasculature, of *runx1-2A-creERT2* dsRed+ HSPCs was similar to the previously described *runx1+23:GFP* labeled anterior mediolateral clusters ^6^. Quantification of dsRed+ cells in the anterior mediolateral clusters showed a steady decrease over 5-7 dpf when 4-OHT induction was performed at later time points, from the end of HSPC DA emergence (50-72 hpf) (Figure 4J – M) and continued and expansion in the CHT (74-96 hpf) (Figure 4N-Q) (Figure 4R-U). The results support Runx1+ HSPCs that colonize the developing kidney from 5 days onward originate from definitive HSPCs in the embryo.

**Figure 4:**
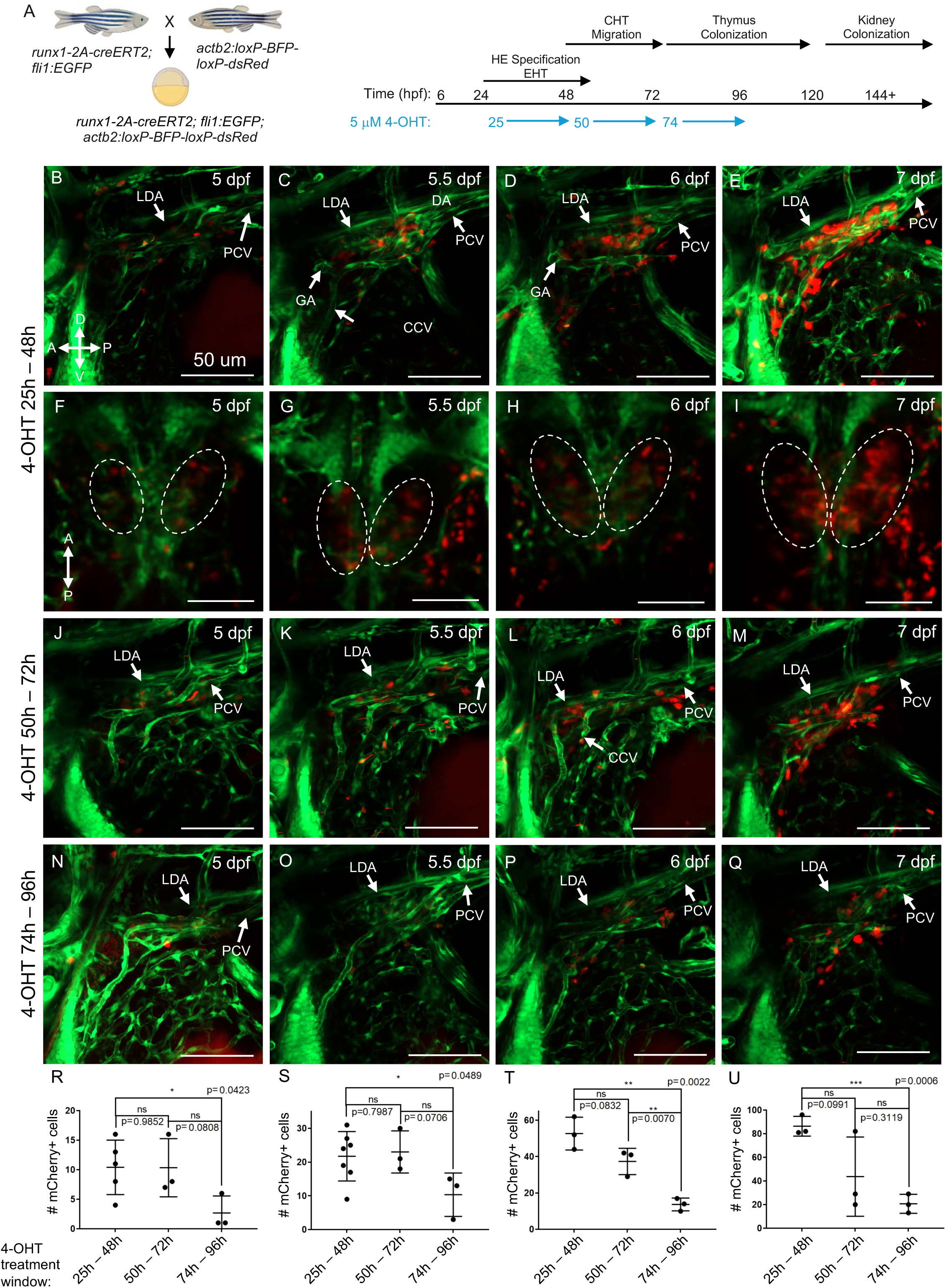
*runx1* lineage tracing in *runx1-2A-creERT2; fli1:EGFP; actb2:loxP-BFP-loxP-dsRed* 5- 7 dpf larva maps kidney colonization from 5+ days post fertilization. A Schematic showing genetic cross to generate *runx1-2A-creERT2; fli1:EGFP; actb2:loxP-BFP-loxP-dsRed* embryos and timeline of 4-OHT treatments during critical events in developmental hematopoiesis. **B –** Q Live confocal imaging of EGFP and dsRed expression in 5, 5.5, 6 and 7 dpf *runx1-2A-creERT2; fli1:EGFP; actb2:loxP-BFP-loxP-dsRed* kidney after 4-OHT treatment from 25-48 hpf (**B-E** lateral, **F-I** dorsal), 50- 72 hpf (**J-M**), and 74-96 hpf (**N-Q**) showing *runx1+* HSPCs in left and right anterior mediolateral clusters (**F-I** dashed circles). **R-U** Quantification of Runx1 dsRed+ HSPCs in the anterior mediolateral clusters at 5 (**R**), 5.5 (**S**), 6 (**T**) and 7 (**U**) dpf after 24-hour window 4-OHT treatments. n=3-7 larva imaged and quantified at each time point. p values calculated using one sample t-test. Plots show mean ± SEM. A, anterior; CCV, common cardinal vein; CHT, caudal hematopoietic tissue; D, dorsal; EHT, endothelial-to-hematopoietic transition; GA, glomerular arteries; HE, hemogenic endothelium; LDA, lateral dorsal aorta; P, posterior; PCV, posterior cardinal vein; V, ventral. Scale bars B-Q 50 µm.

### Lineage tracing developmental and adult Runx1^+^ hematopoiesis

We analyzed the long-term potential of developmentally labeled *runx1* cells to contribute to adult hematopoiesis by inducing lineage tracing with 5 μM 4-OHT from gastrula through 5 dpf, raising the larva to 5 months of age, and performing flow cytometric analysis of kidney marrow and peripheral blood (Figure 5A, Figure S5). 1.2-2.5% of adult kidney marrow cells were mCherry^+^ and present in hematopoietic precursor, myeloid and lymphoid lineages (Figure 5B, Figure S5C). 1.2-4.9% of peripheral blood cells, mainly composed of erythrocytes, were mCherry^+^ (Figure 5C, Figure S5C). Lineage labeling was also analyzed in 3 individual adults by treatment of 7-month-old *runx1-2A- creERT2; ubi:Switch* with 5 μM 4-OHT for 12 hours per day over three days, followed by analysis on day 9 (Figure 5D). Adult lineage labeling resulted in 8-10% mCherry^+^ kidney marrow cells, present in precursor, myeloid and lymphoid lineages (Figure 5E, Figure S5C), and 0.1-2.5% mCherry^+^ peripheral blood cells (Figure 5F, Figure S5C). Flow cytometry analysis of kidney marrow or peripheral blood from adults treated with EtOH vehicle alone, either as embryos (Figure 5B, C) or adults (Figure 5E, F) was negative for mCherry^+^ cells. Runx1 labeling of adult hematopoietic cells was consistent with single cell gene expression analysis of *Runx1* human ^47^ and r*unx1* zebrafish ^36^ in adult marrow lineages (Figure S6). Together, these results demonstrate zebrafish Runx1 embryonic progenitors give rise to a population of long lived hematopoietic precursors that persist into adulthood and give rise to all adult blood lineages.

**Figure 5:**
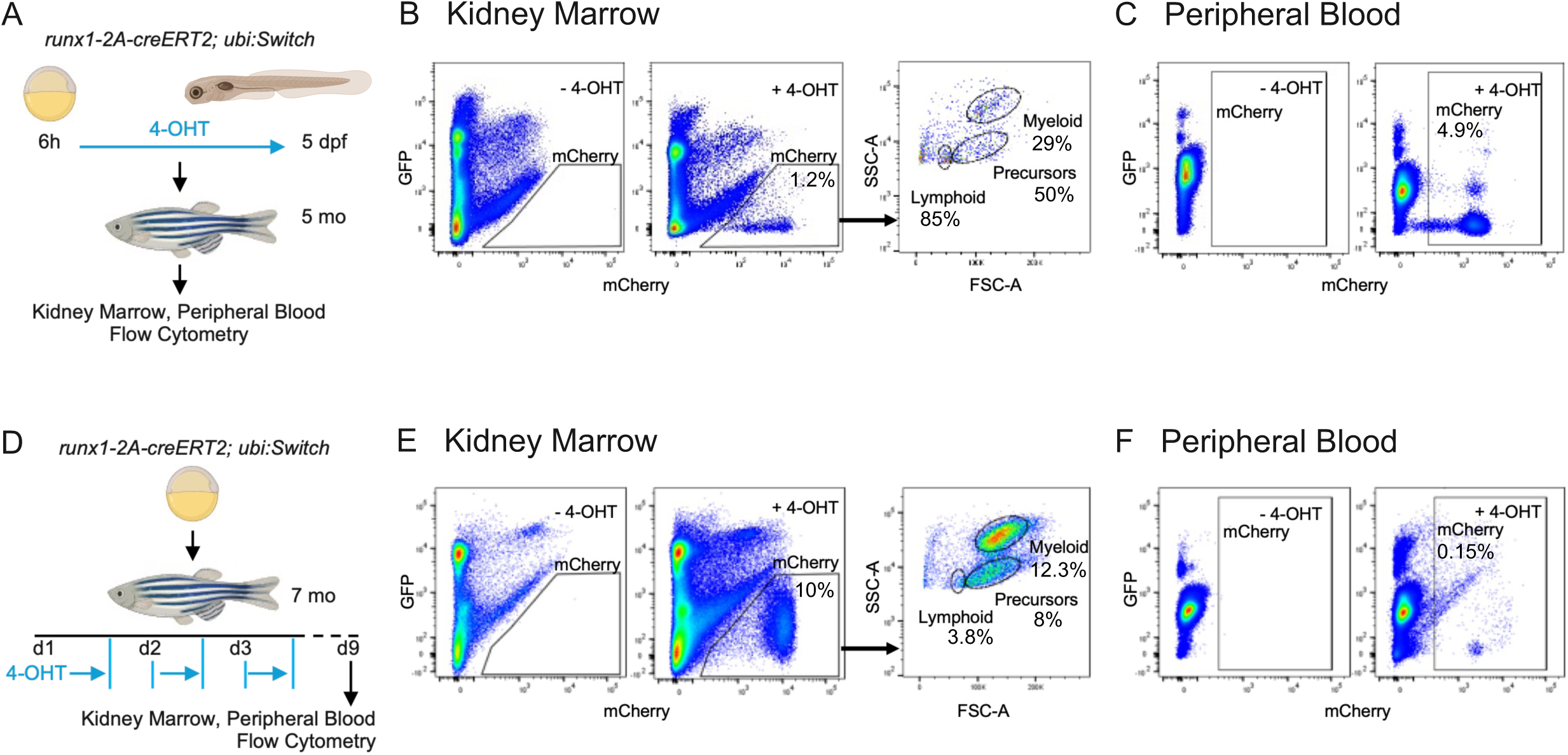
Long-term and adult lineage tracing of *runx1^+^* cells in adult kidney marrow and peripheral blood. **A–C** Long-term lineage tracing. n=5 adult fish analyzed per treatment. **A** Schematic showing 4-OHT treatment of *runx1-2A-creERT2; ubi:Switch* embryos from shield stage (6h) through 5 dpf, followed by kidney marrow and peripheral blood flow cytometric analysis at 5 months. **B** Representative flow cytometric analysis of mCherry^+^ cells in kidney marrow from embryo switched *runx1-2A-creERT2; ubi:Switch* adults (+4-OHT). Flow cytometric light scatter profile of kidney marrow mCherry^+^ cells showing Precursor, Myeloid and Lymphoid hematopoietic lineages. **C** Representative flow cytometric analysis of mCherry^+^ peripheral blood from embryo switched *runx1- 2A-creERT2; ubi:Switch* adults (+4-OHT). **D – F** Adult lineage tracing. n=3 adult fish analyzed per treatment. **D** Schematic showing 4-OHT treatment of 7-month *runx1-2A-creERT2; ubi:Switch* adults for 12 hours per day for three days, followed by kidney marrow and peripheral blood flow at 6 days post treatment. **E** Representative flow cytometric analysis of mCherry^+^ cells in kidney marrow from *runx1-2A-creERT2; ubi:Switch* switched adults (+4-OHT). Flow cytometric light scatter profile of mCherry^+^ cells showing Precursor, Myeloid and Lymphoid hematopoietic lineages. **F** Representative flow cytometric analysis of mCherry^+^ peripheral blood from *runx1-2A-creERT2; ubi:Switch* switched adults (+4-OHT). Control plots (-4-OHT) show results from ethanol treatment alone.

## Discussion

In this study we describe a zebrafish *runx1-2A-CreERT2* precision knock-in line and demonstrate its tamoxifen-regulated Cre recombinase activity recapitulates the complete *runx1* lineage of neuromesodermal and lateral plate mesodermal derived progenitors. The *2A-creERT2* cassette was integrated at the 3’ end of zebrafish *runx1* exon 7, enabling Cre expression from all reported zebrafish *runx1* promoters and mRNA isoforms ^2,3^. We traced the lineage of *runx1* hematopoietic cells from the embryo hemogenic endothelium to larval hematopoietic organs and the adult kidney marrow, and demonstrated embryonic *runx1* hematopoietic cells persist to adulthood. Our results underscore the advantage of endogenous Cre knock-ins, where the pattern of Cre expression is precisely controlled by the complete landscape of gene regulatory elements, which can be difficult to achieve even with large BAC genomic DNA promoter:reporter transgenics ^2^. Zebrafish *runx1-2A-CreERT2* provides the ability to perform inducible, spatial and temporal analysis of Runx1 lineages and HSPCs without ectopic or leaky recombinase activity, in development and in adult stages.

Tracing *runx1-2A-CreERT2* progenitors in embryonic blood lineages labeled both primitive and definitive hematopoietic cells. Time lapse analysis of the embryo DA showed *runx1*-labeled definitive HSPCs budding from the DA ventral wall and entering circulation in the PCV, confirming that *runx1* lineage tracing identifies the zebrafish equivalent of the mammalian AGM. *runx1*-negative cells were also detected budding from the DA. This may reflect lower levels of Switch reporter mCherry transgene expression in some cells, or represent the recently described Runx1-independent DA lineage that gives rise to vessel resident macrophages ^48^.

The *runx1-2A-creERT2* line mapped definitive Runx1+ HSPCs to the caudal hematopoietic tissue, thymus and pronephros larval hematopoietic organs. Previous studies in zebrafish have characterized HSPC DA emergence and mapping to hematopoietic organs using endothelial and stem cell gene fluorescent reporters and photoactivatable cell tracers ^1,6,7,23,24,26,46,49^. Our results are consistent with these analyses, and further demonstrate using timed induction of Cre activity that Runx1 progenitors colonize the larval thymus and kidney during the HSPC expansion stage in the caudal hematopoietic tissue. CreERT2 induced tracer switching during DA emergence from 24 – 50 hpf mapped Runx1 precursors to the thymus and anterior pronephros, while later switching post 3 dpf significantly reduced Runx1 labeled cells in both organs. The reduction in *runx1-2A-creERT2* thymus at later induction time points at 5 dpf is consistent with previous observations showing an absence of thymic *runx1* expression ^24^ and supports downregulation of *runx1* in lymphoid fated precursors destined for migration to the thymus. In contrast to thymic precursors, cells in the glomerulus region of the 5+dpf larva express *runx1* ^24^. Our results show *runx1-2A-creERT2* labeled HSPCs populate the anterior mediolateral clusters at 5+ dpf, providing support for Runx1 precursors colonizing the larval pronephros and for our demonstration of long term lineage tracing of *runx1*-derived hematopoietic precursors and blood cells in adult kidney and peripheral blood. It is possible *runx1*-labeled thymic cells originated from a population of HSC-independent lymphoid precursors ^7,45^. Previous analyses reported two independent CHT-resident progenitor populations give rise to embryonic blood or to definitive HSCs that support adult hematopoiesis ^7^. While our results using temporal induction of CreERT2 labeling support Runx1 precursors derived from embryonic definitive hematopoiesis populate the future adult kidney, additional studies with the *runx1-2A-creERT2* line could further define the developmental origin of definitive and functional HSPCs.

Long-term lineage tracing demonstrated *runx1-2A-creERT2* labeled embryonic HSPCs persisted into adulthood and contributed to all blood lineages. Previous analysis with a broad vascular endothelial *kdrl:Cre* driver demonstrated zebrafish HSCs derive from the embryo aortic endothelium ^23^. In contrast, the *runx1-2A-creERT2* line described here restricts hematopoietic lineage tracing specifically to aortic hemogenic endothelium, and allows temporal induction to separate developmental and adult hematopoiesis. *runx1-2A-creERT2* lineage tracing induced in adult fish labeled all hematopoietic lineages, confirming Runx1 activity throughout the adult hematopoiesis system, consistent with prior reports using the *runx1+23:GFP* transgenic line ^36^. Early mouse studies reported Runx1 is broadly expressed in adult HSCs, myeloid and lymphoid cells ^37^, while more recent single cell analysis in mouse ^47^ and zebrafish ^36^ indicate relatively higher levels in myeloid lineages. Functional studies are needed to determine whether Runx1 progenitors are programmed to differentiate into adult blood cells with a potential myeloid bias.

The zebrafish *runx1-2A-creERT2* line enables tamoxifen-controlled spatial and temporal Runx1 lineage analysis. In combination with Cre/*lox* regulated genes, *runx1-2A-creERT2* provides a powerful tool to investigate genetic mechanisms of hematopoiesis, such as further defining the requirement for and compensatory role of *gata2* in blood development ^50,51^. The ability to now perform robust HSPC- specific, cell autonomous gene studies, as we demonstrated previously in proneural progenitors in the developing nervous system ^12^, will enable new conditional gene models of human blood disorders and inflammatory disease.

Our results are consistent with the long-standing view that HSPCs specified during embryonic development persist throughout life and give rise to adult blood, and show *runx1* embryonic progenitors contribute to hematopoietic precursors in zebrafish adult kidney marrow. Recent studies describe a new HE HSPC niche in the zebrafish larval supra-intestinal artery (Feng et al., 2025 bioRxiv **doi:** https://doi.org/10.1101/2025.05.20.655066), and HSC-independent progenitors in the mouse embryo ^52–54^ and adult ^55^ hematopoietic compartments. Whether Runx1 progenitors contribute to these novel hematopoietic populations is an outstanding question that can be addressed using *runx1-2A-creERT2* genetic analysis.

## Limitations

Zebrafish CreERT2 recombinase lines generated by integration of *creERT2* into endogenous genes, such as the *runx1-2A-creERT2* precision knock-in described here, provide important advantages for experimental reproducibility by overcoming limitations of traditional transgenesis that can result in overexpression, ectopic expression, and leaky Cre recombinase activity due to copy number variation and position effect. Consequently, the activity of traditional zebrafish *runx1* transgenics is not comparable to the endogenous *runx1-2A-creERT2* knock-in line described here. *runx1-2A-creERT2* traces all expected developmental HSPC populations, and all adult hematopoietic lineages after embryonic and adult 4-OHT induction. The protocol for *runx1-2A-creERT2* 4-OHT induction in developmental and adult stages can be further optimized for specific life stages and tissue analyses.

## Materials and Methods

### Zebrafish Care and Husbandry

Zebrafish (*Danio rerio*) were maintained on an Aquaneering aquaculture system at 27°C on a 14 hour light/10 hour dark cycle. The *runx1* Knock-in line described here was generated in the wild type WIK background. WIK adults were obtained from the Zebrafish International Resource Center (https://zebrafish.org/home/guide.php). Previously reported transgenic reporter lines used in this study: *Tg(-3.5ubb:loxP-EGFP-loxP-mCherry)cz1701* ^43^ (*ubi:Switch*), *Tg(actb2:loxP-BFP-loxP- dsRed)sd27* ^23^, and *Tg(fli1:EGFP)y1* ^56^ (*fli1:EGFP*). Wildtype and transgenic embryos were collected and maintained at 28.5°C in E3 embryo media ^57^ and staged according to published guidelines ^58^.

### Ethics approval statement

Experimental protocols used in this study were approved by the Iowa State University Institutional Animal Care and Use Committee (IACUC-20-058, IACUC-20-025, IBC-20-071, IBC-20-024), in compliance with American Veterinary Medical Association and the National Institutes of Health guidelines for the humane use of laboratory animals in research. Experiments reported in this study follow the ARRIVE guidelines ^59,60^.

### Isolation of *runx1-2A-creERT2, exorh:CFP^is97^* with GeneWeld CRISPR-Cas9 targeted integration

The *2A-creERT2, exorh:CFP* cassette was integrated in frame at a CRISPR-Cas9 cut site located 29 nucleotides upstream of the TGA translation stop in *runx1* exon 7, using GeneWeld targeted integration, as described previously ^9,10^. The sequences of gRNA and oligonucleotides used in this study are shown in S1. A reverse strand synthetic guide targeting zebrafish *runx1* (ZDB-GENE- 000605-1) 5’-GACGGCCTCTACAGACATGC-3’ (Integrated DNA Technologies) was tested for mutagenesis by co-injection of 5 pg gRNA with 150 pg *in vitro* transcribed, capped, polyadenylated nlsCas9nls mRNA, into single cell zebrafish embryos. The *runx1* target site was amplified with *runx1* exon 7 F and R primers (Table S1), amplicons Sanger sequenced, and mutagenesis efficiency analyzed using ICE software (Synthego). The *gcry:eGFP* secondary reporter in the GeneWeld *pPRISM-2A-cre, gcry:eGFP* construct ^10^ was replaced with a pineal:Cerulean expression cassette *exorh:CFP* ^40^ using New England BioLab HiFi cloning (NEB #E2621S). *runx1* target site 24 bp 5’ homology arm with mutated PAM C**C**A → C**A**A, and 48 bp 3’ homology arm, were cloned into the *pPRISM-2A-cre, exorh:CFP* construct. The final *runx1-2A-creERT2, exorh:CFP* targeting vector was assembled by replacing the *cre* cDNA with *creERT2* from *pENTR_D_creERT2* ^43^ (Addgene 27321) as described ^11^ and confirmed by whole plasmid sequencing (Plasmidsaurus). From 24 adult fish screened, nine founders transmitted the pineal-specific Cerulean marker, and two of the nine contained a precise in frame integration of *runx1-2A-creERT2, exorh:CFP* at the genomic target site. One founder transmitting a precise allele was outcrossed to wild type, and a single F1 adult confirmed by fin clip and sequencing, was used to establish the knock-in line *runx1-2A-creERT2, exorh:CFP^*is*97^*.

### RNASeq analysis

*runx1creERT2* F3 zebrafish were incrossed to produce +/+, *runx1-2A-creERT2*/+ and *runx1-2A- creERT2/runx1-2A-creERT2* embryos. Embryos were raised to 28 hpf in E3 embryo media. Individual embryos were deyolked for genotyping and flash frozen at -80. Ten genotyped embryos were pooled (n=3 biological replicates for each genotype) and total RNA extracted from each pool using the RNeasy Mini Kit (Qiagen 74104) and purified by RNA Clean & Concentrator™-5 (Zymo Research R1013). Bulk RNASeq oligo-dT primed first strand cDNA synthesis, library preparation and Illumina sequencing were performed at Plasmidsaurus. Gene enrichment analysis and Volcano plots were provided by Plasmidsaurus. Additional differential gene expression analysis was performed using iDEP 2.0 (https://bioinformatics.sdstate.edu/idep20/) and heat maps were created in Heatmapper (https://heatmapper.ca). RNASeq raw data deposited at GEO (submission in progress).

### Whole-mount RNA *in situ* hybridization (WISH), Quantification and Statistical Analysis

WISH was performed as described previously ^61^. Heterozygous *runx1-2A-creERT2*-/+ adults were in- crossed and all embryos collected and fixed in 4% paraformaldehyde at 28 hpf and 42 hpf. *runx1* and *cmyb* probes for *in situ* were generated from linearized pCS2 plasmid templates using the DIG RNA Labeling Kit (MilliporeSigma; 11175025910)^62^. Embryos were imaged in PBS using a Leica M165FC stereomicroscope equipped with a DFC295 color digital camera (Leica) and FireCam software (Leica) with Leica Application Suite X (v3.7.0.20979). *runx1* and *cmyb* WISH labeled dorsal aorta cells were quantified in a double blind experiment. After quantification the genotype of each embryo was determined by genomic DNA PCR ^9^ with *runx1* exon 7 F, runx1 exon 7 R, and 5’-junction-R knock-in cassette primers listed in Table S1. Statistical analyses were performed with GraphPad Prism Software using unpaired Student’s t-test.

### Tamoxifen treatment and imaging

Analysis of the complete *runx1* progenitor lineage derived was performed by treatment of *runx1-2A- creERT2*; *ubi:Switch* embryos continuously from shield stage through 5 days post fertilization (dpf) with 5 μM 4-hydroxytamoxifen (4-OHT) (Sigma H6278) in E3 embryo media. For temporal analysis of *runx1* progenitor contribution to definitive hematopoiesis, *runx1-2A-creERT2*; *ubi:Switch* and *runx1- 2A-creERT2*; *fli1b:EGFP*; *actb2:loxP-BFP-loxP-dsRed* embryos or larvae were treated with 5 μM 4- OHT at four time intervals: from 6 hpf to 24 hpf, 26 hpf to 48 hpf, 50 hpf to 72 hpf, and 74 hpf to 96 hpf. Following exposure to 4-OHT the embryos or larvae were placed in fresh E3 media until imaging. Control embryos or larvae were treated with 0.05% EtOH vehicle alone. For live imaging embryos were incubated in E3 embryo media ^57^ containing 0.003% 1-phenyl 2-thiourea (PTU) (Sigma P7629) to inhibit pigment synthesis. Embryos and larvae were anesthetized in 0.015% Tricaine MS-222 Ethyl 3-aminobenzoate methanesulfonate (Sigma E10521) and mounted in 1.2% low melt agarose (Promega V2111) in embryo media containing 0.003% PTU/0.015% Tricaine. Live embryo and larvae fluorescence and differential interference contrast images were captured using a Zeiss LSM 800 laser scanning confocal microscope. Statistical analyses were performed with GraphPad Prism Software using one sample t-test.

### Kidney dissection and flow cytometric analysis of kidney marrow and peripheral blood

Long term tracing of *runx1* HSPC lineages in the adult kidney was performed by treating *runx1-2A- creERT2*; *ubi:Switch* embryos with vehicle alone or 5 μM 4-OHT from 6 hpf to 5 dpf and raising adults to 5 months. Lineage labeling of *runx1* hematopoietic cells in adult kidney was performed by treatment of 7-month-old *runx1-2A-creERT2*; *ubi:Switch* adults with 5 μM 4-OHT in 200 ml fish water, or ethanol vehicle alone, for 12 hours per day for 3 days, and sacrifice 6 days later for flow cytometry. Adult zebrafish were anesthetized in 0.2% Tricaine MS-222 Ethyl 3-aminobenzoate methanesulfonate (Sigma E10521), peripheral blood collected, kidney dissected, and dissociated cells analyzed by flow cytometry as previously described ^63,64^. Kidney marrow suspension was filtered through a 30µm cell strainer (Thermo Fisher Scientific, NC9084441) and stained with SYTOX™ Red (Life Technologies; S34859) to exclude dead cells. Flow cytometric acquisitions were performed on a FACS Melody (BD) with FACSChorus 2.0, version 1.1.20.0 software, and analyses using FlowJo software (v10.3 or v10.8.1, Tree Star) ^63^. Flow analysis on adult kidney marrow and peripheral blood was performed in triplicate using three adult individuals for each condition (vehicle treated control *runx1-2A-creERT2*; *ubi:Switch*, 4-OHT treated *runx1-2A-creERT2*; *ubi:Switch*).

## Supporting information

Detailed Protocol

## Acknowledgements

The authors thank the Iowa State University DNA Facility (Office of Biotechnology) for performing Sanger sequencing. The *exorh* gene promoter:Cerulean Fluorescent Protein cDNA cassette *exorh:CFP* was a gift from Dr. Christian Mosimann, U Colorado Denver Anschutz Medical Campus. The authors thank Dr. Fang Liu, Iowa State University, Dr. D’Juan Farmer, UCLA, Dr. Owen Tamplin, University of Wisconsin-Madison and Dr. Teresa Bowman, Albert Einstein College of Medicine for discussions.

## Author Contributions

Conceptualization: JAP, RE-P, MM

Methodology: JAP, MKU, RE-P, MM

Investigation: JAP, MKU

Visualization: JAP, MKU, RE-P, MM

Funding acquisition: MM, RE-P, JJE, KJC, SCE

Project administration: MM

Supervision: MM, RE-P

Writing – original draft: JAP, MM

Writing – review & editing: JAP, KJC, SCE, JJE, RE-P, MM

## Declaration of Interests

JJE is a shareholder/founder of Immusoft Inc., LifEngine and LifEngine Animal Health.

KJC is a shareholder/founder of LifEngine and LifEngine Animal Health

SCE is a shareholder/founder of LifEngine and LifEngine Animal Health

MM is an immediate family member of JJE.

JJE, KJC, SCE, MM disclose US Patent 12,305,168 B2. Materials and methods for efficient targeted knock in or gene replacement. Inventors Essner, J.J., Clark, K.J., Ekker, S.C. McGrail, M.A., Wierson, W.A., Welker, J., de Almeida, M.P. May 20, 2025. Rochester, MN. Ames, IA.

JAP, MKU, RE-P declare no competing interests.

## Data Availability

DNA constructs and *runx1-2A-creERT2, exorh:CFP^is97^* zebrafish line are available on request to M. McGrail.

## Supplementary Information

**Document S1.** Supplemental Figures S1–S5, Table S1.

**Table S2, related to Figure 1**. Bulk RNASeq wildtype, *runx1-2A-creERT2/+* heterozygous, *runx1-2A- creERT2/runx1-2A-creERT2* homozygous 28 hpf embryos. **Separate excel file**

**Video S1, related to Figure 2**. Confocal time lapse video of Runx1-mCherry^+^ HSPCs during emergence from the embryo DA at 30 hpf. **.mp4 file**

**Video S2, related to Figure 2**. Confocal time lapse video of Runx1-mCherry^+^ HSPC during emergence from the embryo DA at 48 hpf. **.mp4 file**

**Video S3, related to Figure 2**. Confocal time lapse video of Runx1-mCherry^+^ precursors in the larval thymus at 4 dpf. **.mp4 file**

**Video S4, related to Figure 2**. Confocal time lapse video of Runx1-mCherry^+^ precursors in the larval thymus at 5 dpf. **.mp4 file**

**Detailed Protocol.** Protocols for tamoxifen regulated CreERT2 tracing of Runx1 lineages in zebrafish *runx1-2A-CreERT2* embryos, larvae and adults. **Separate pdf file**

