## Supplemental Figures and Table S1 for "Tracing developmental and adult hematopoiesis with an endogenous zebrafish *runx1-2A- CreERT2* CRISPR knock-in"

**Document S1.** Supplemental Figures S1–S5, Table S1.

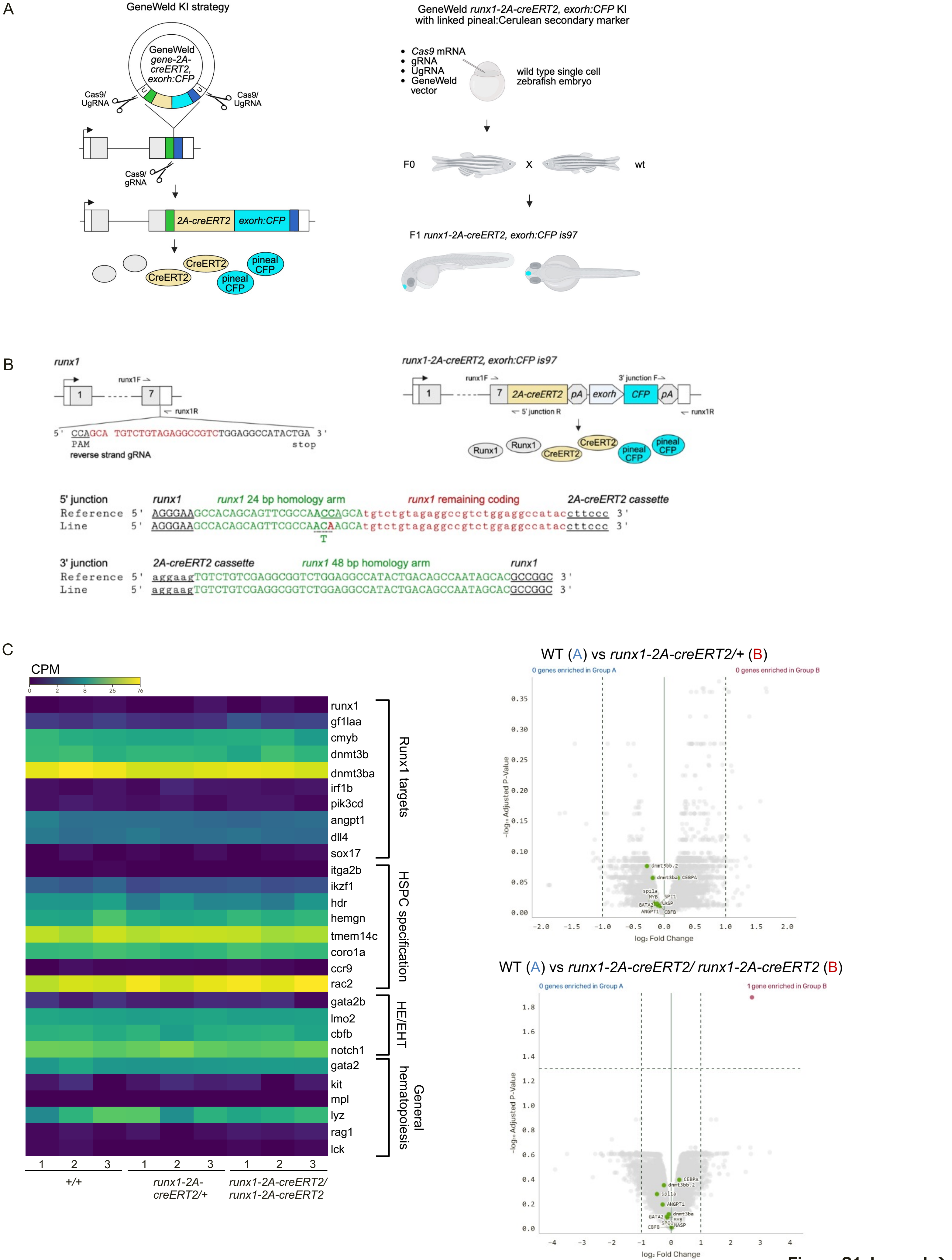


**Figure S1 Legend →**

**Figure S1, related to Figure 1. *runx1-2A-creERT2* knock-in line generated with GeneWeld CRISPR-Cas9 targeted integration. A** GeneWeld strategy for targeted integration of *2A-creERT2,exorh:CFP* cassette into 3′ end of gene coding sequence. Diagram of experimental workflow for injection and line recovery. **B** Left, *runx1* diagram showing sequence of exon 7 gRNA in red. Space, Cas9 cut site. PAM in bold and underlined. Primers for generating PCR amplicon to test *runx1* gRNA mutagenesis efficiency are labeled runx1F and runx1R. Right, diagram of recovered *runx1-2A-creERT2,exorh:CFPis97* line with cassette integrated in frame at the 3′ end of the coding sequence. Location of primers used for 5′ and 3′ junction PCR analysis. Sanger sequence results of 5′ and 3′ genomic DNA-cassette knock-in junction PCR amplicons from *runx1-2A-creERT2,exorh:CFPis97* line. 24 bp 5′ and 48 bp 3′ homology arms are shown in green. 5' homology arm silent C>A mutation in PAM shown in red. Dashed underline beneath Threonine codon (T). Sequences added to the 5′ homology arm to complete the *runx1* coding sequence shown in red lowercase. **C** Bulk RNAseq heat map and volcano plot comparisons of differential gene expression in 28 hpf wildtype, *runx1-2A-creERT2/*+ heterozygotes and *runx1-2A-creERT2/ runx1-2A-creERT2* homozygotes. CPM, counts per million; HE, hemogenic endothelium; EHT, endothelial to hematopoietic transition.

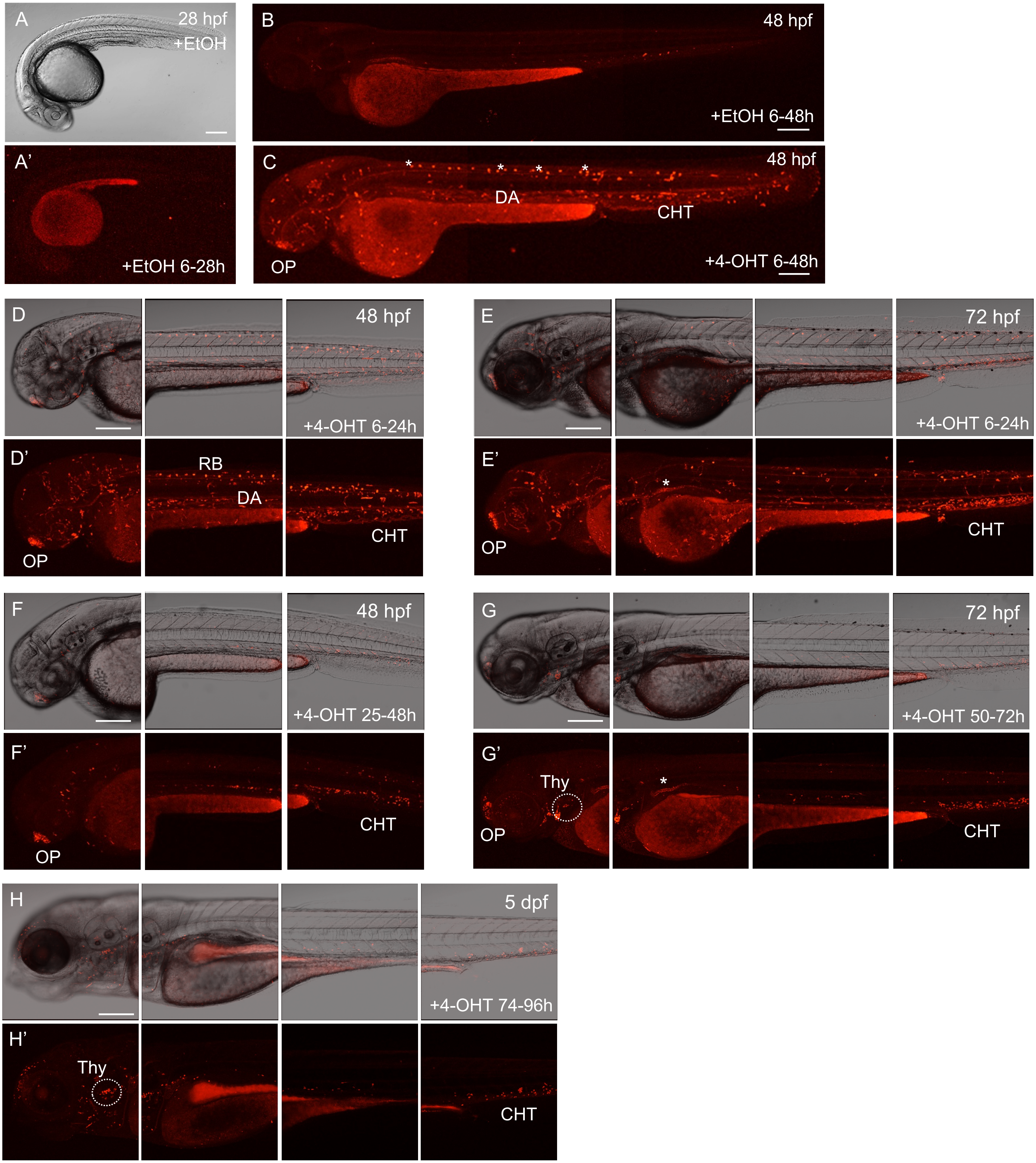


**Figure S2, related to Figure 2 and Figure 3. *runx1-2A-creERT2; ubi:Switch* lineage tracing whole embryo and larva imaging after 24 hour 4-OHT treatment windows.** Live confocal imaging of mCherry expression in *runx1-2A-creERT2; ubi:Switch* embryos and larvae. EtOH control treated 28 hpf (**A, A'**) and 48 hpf (**B**) embryos. 48 hpf embryo treated with 4-OHT from 6-48 hpf (**C**), asterisks mark RB neurons. Embryos and larvae after 4-OHT treatment from 6-24 hpf (**D, D', E, E'**), 25-48 hpf (**F, F'**), 50-72 hpf (**G, G'**), and 74-96 hpf (**H, H'**). Asterisks in E' and G' mark non-specific signal in pronephros anterior proximal tubule. DA, dorsal aorta; OP, olfactory placode; RB, Rohan-Beard neurons; Thy, thymus. n=3 larva imaged at each time point. Scale bars A – H 200 μm.

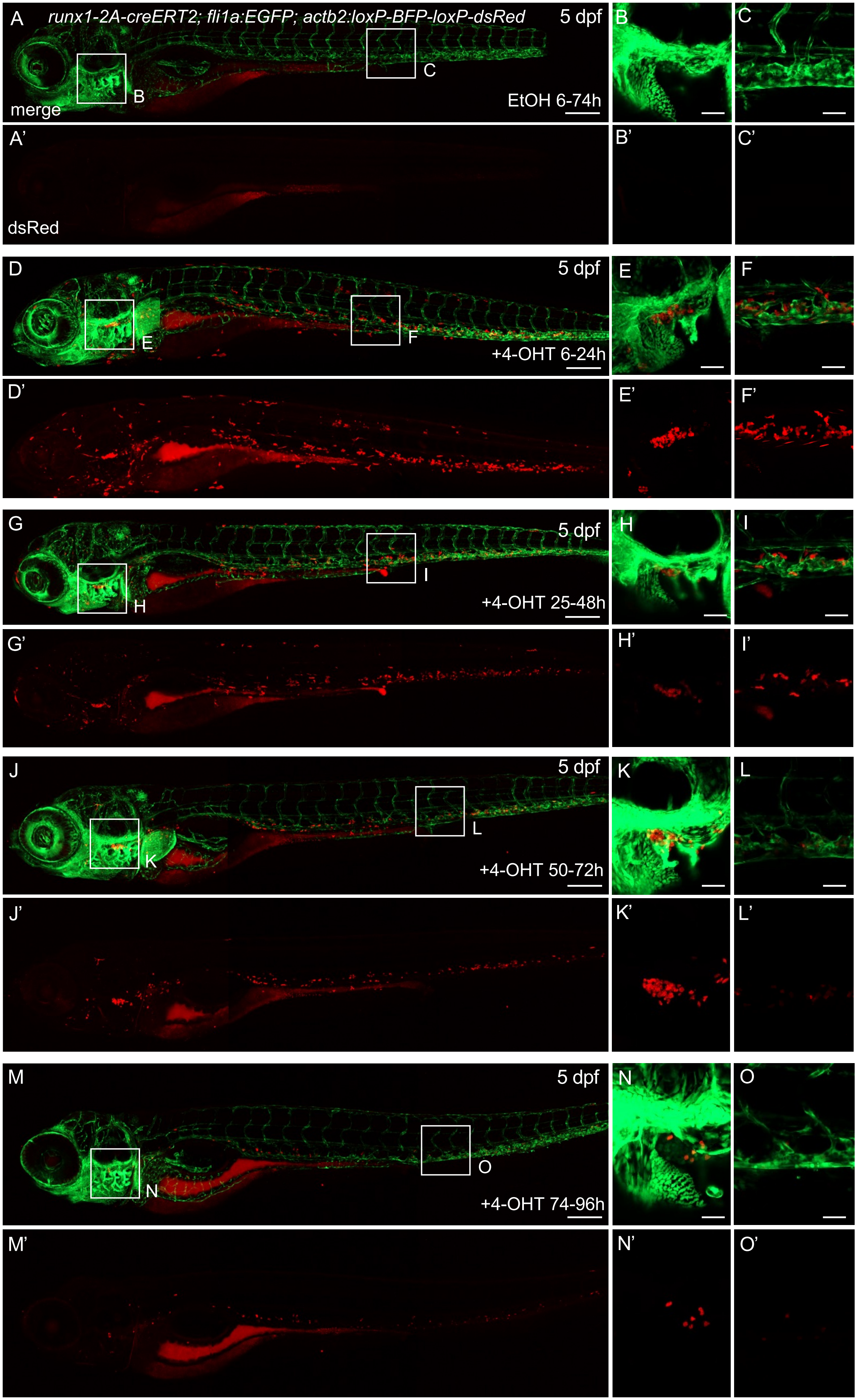


**Figure S3 legend →**

**Figure S3, related to Figure 4.** ***runx1*** **lineage tracing in** ***runx1-2A-creERT2; fli1a:GFP;actb2:loxP-BFP-loxP-dsRed*** **larva after 24 hour 4-OHT treatment windows beginning at shield stage, 25 hpf, 50 hpf, and 74 hpf.** Live confocal imaging of EGFP and dsRed expression in 5 dpf *runx1-2A-creERT2; fli1a:GFP;actb2:loxP-BFP-loxP-dsRed* larvae. Control EtOH treatment from 6-74h (**A, A', B, B', C, C'**) or 4-OHT treatment from 6-24 hpf (**D, D', E, E', F, F'**), 25-48 hpf (**G, G', H, H', I, I'**), 50-72 hpf (**J, J', K, K', L, L'**), and 74-96 hpf (**M, M', N, N', O, O'**). n=3 larva imaged for each treatment condition. Scale bars A,D,G,J,M 200 µm; B,C,E,F,H, I,K,L,N,O 50 µm.

**A**

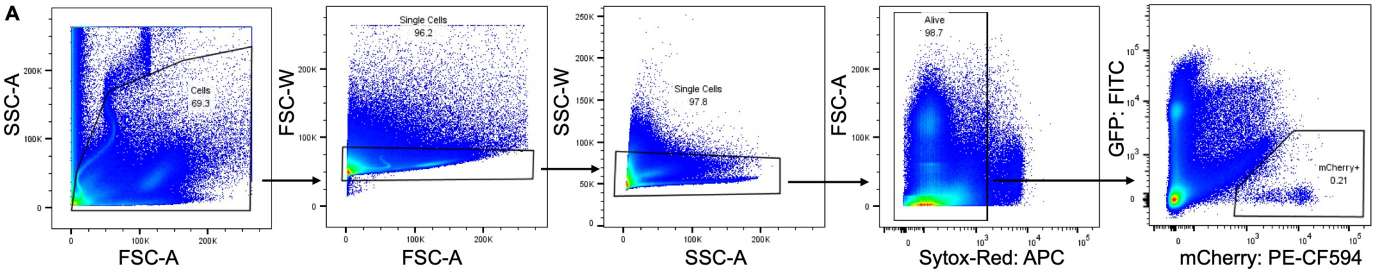


**B**

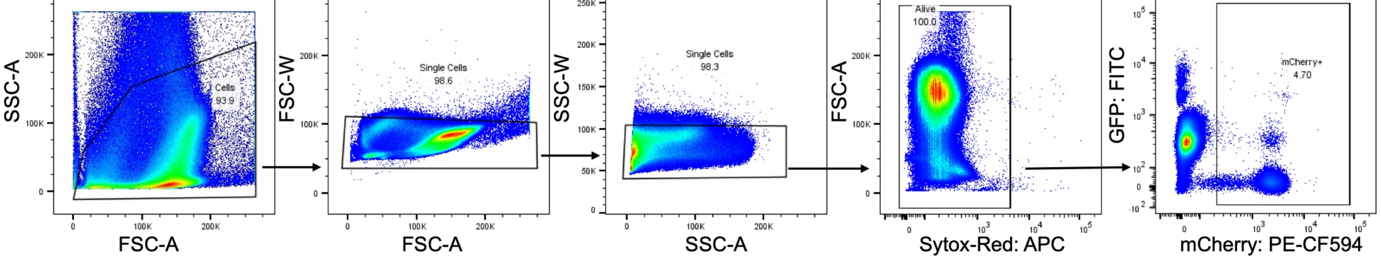


**C**

| % mCherry+ cells | | | | |
|---|---|---|---|---|
| Replicate | Long term, kidney marrow | Long term, peripheral blood | Adult treatment, kidney marrow | Adult treatment, peripheral blood |
| 1 | 1.16 % | 2.31 % | 10.26 % | 0.147 % |
| 2 | 1.19 % | 1.38 % | 8.32 % | 2.53 % |
| 3 | 2.44 % | 4.86 % | 1.22 % | 0.06 % |
| 4 | 2.26 % | 2.53 % | - | - |
| 5 | 1.59 % | 1.24 % | - | - |

**Figure S4, related to Figure 5. Gating strategies and flow analysis of runx1-2A-creERT2 lineage tracing in adult marrow and peripheral blood. A** Representative gating strategies from adult kidney marrow flow cytometric data in Figure 4B, E. **B** Representative gating strategies from peripheral blood flow cytometric data in Figure 4C, F. **C** Table showing percentages of mCherry+ cells identified by flow cytometry in adult kidney marrow and peripheral blood after 4-OHT induction during embryogenesis (Long term) and in adults (Adult treatment).

A

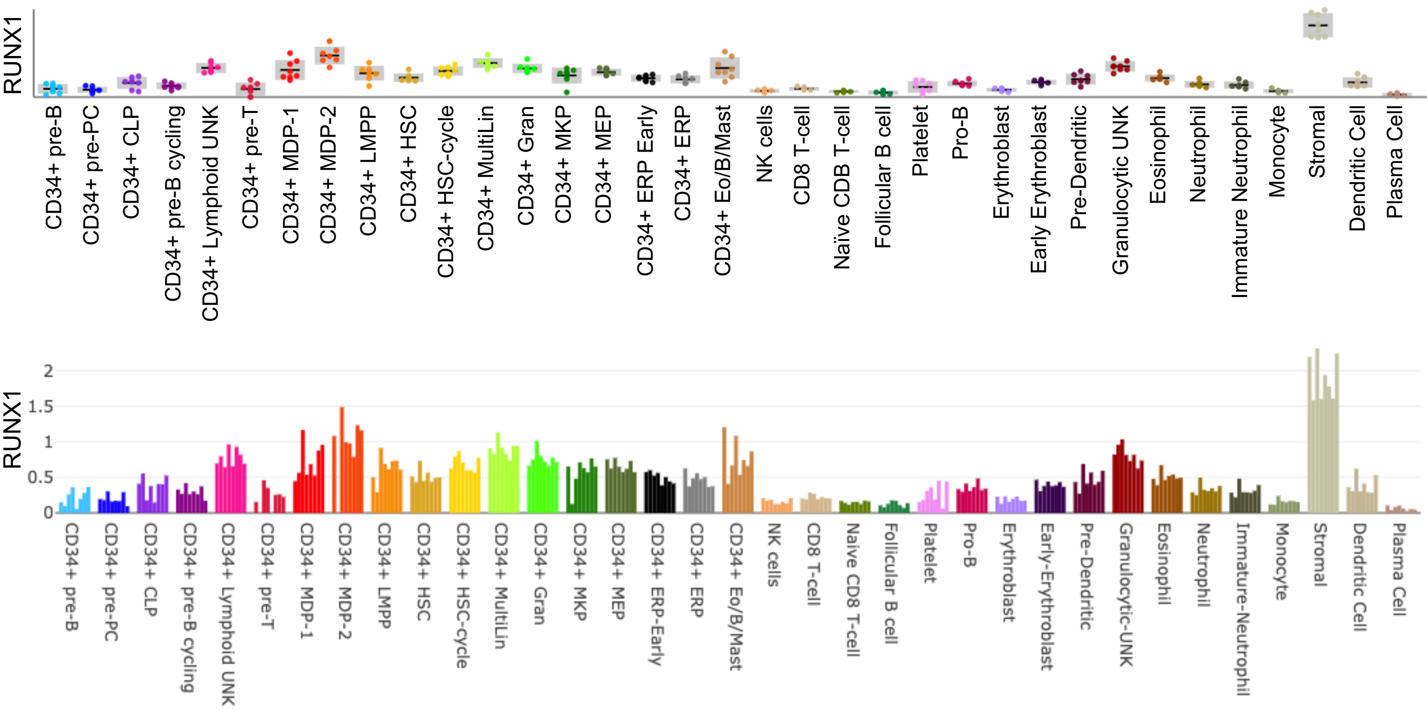


B

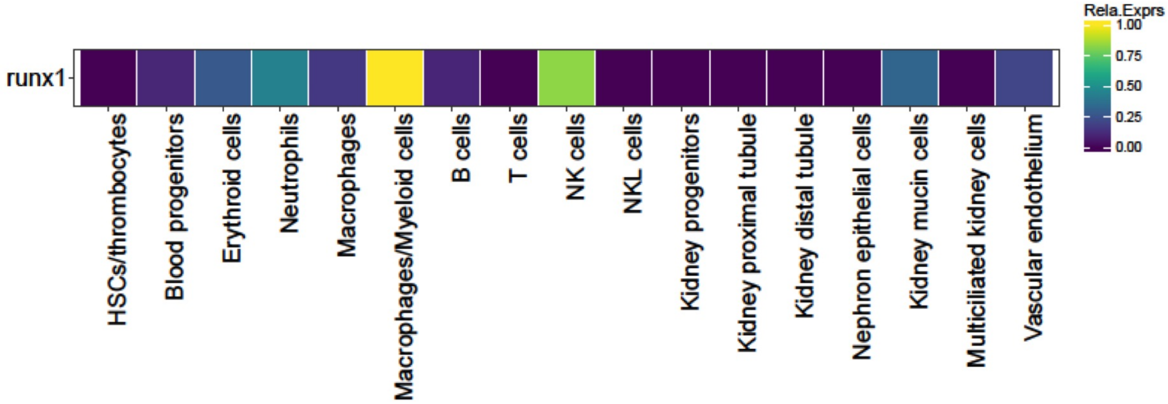


**Figure S5, related to Figure 5. RUNX1 is expressed in all hematopoietic lineages in human bone and zebrafish kidney marrows. A** Box plot and bar plot data extracted from the Human bone marrow cell atlas portal (https://altanalyze.org/MarrowAtlas/human.html https://www.altanalyze.org/ICGS/HCA/splash.php) (Hay et al., 2018) showing *RUNX1* expression in human bone marrow. Y axis, mean gene expression for all 35 cell-populations for each donor (counts per 10,000 normalized). **B** Heat map showing *runx1* expression based on single-cell RNA-seq data from zebrafish adult kidney marrow generated using the online visualizer “Single Cell inDrops RNA-Seq Visualization of Adult Zebrafish Whole Kidney Marrow” (https://molpath.shinyapps.io/zebrafishblood/#pltly) (Tang et al., 2017.

**Table S1. Sequences of oligonucleotides used in this study.** Oligonucleotide sequences of *runx1* target site gRNA, primers used for PCR, and oligonucleotides to build *runx1* 48 bp homology arm *pPRISM -2A-creERT2, exorh:CFP* targeting vector. Sequences in lowercase represent the remainder of the runx1 coding sequence before the TGA stop codon. Sequences in lowercase italics are sequences complementary to BfuAI and BspQI overhang after restriction enzyme digestion of the GeneWeld *pPRISM-2A-cre* vector. *runx1* 5' homology arm oligonucleotides 1 and 2 contain a C>**A** mutation (bold, underlined) in the PAM to prevent Cas9/CRISPR gRNA cutting.

| **Oligonucleotides** | **Sequence 5' – 3'** | **Purpose** |
|---|---|---|
| *runx1* gRNA | GACGGCCTCTACAGACATGCTGG | exon 7 gRNA with PAM for CRISPR-Cas9 targeting |
| *runx1* exon 7 F | CAGTTCTCCATGATGCCCAGC | PCR amplicon, genome/vector PCR 5' junction analysis |
| *runx1* exon 7 R | CAAAATAAAGTCTAAAGCATGTCAAATTAATTGACTTTTATG | PCR amplicon, genome/vector PCR 3' junction analysis |
| *runx1* 5' homology arm oligo 1 | *gcgg*GCCACAGCAGTTCGCCAAC**A**AGCAtgtctgtagaggccgtctggaggccatac | Clone 5' 24 bp homology arm plus remaining 29 bp of *runx1* coding sequence in GeneWeld vector for CRISPR-Cas9 targeted integration |
| *runx1* 5' homology arm oligo 2 | *gaag*gtatggcctccagacggcctcatcagacaTGCT**T**GTTGGCGAACTGCTGTGGC | Clone 5' 24 bp homology arm plus remaining 29 bp of *runx1* coding sequence in GeneWeld vector for CRISPR-Cas9 targeted integration |
| *runx1* 3' homology arm oligo 1 | *aag*TGTCTGTCGAGGCGGTCTGGAGGCCATACTGACAGCCAATAGCAC | Clone 3' 48 bp homology arm in GeneWeld vector for CRISPR-Cas9 targeted integration |
| *runx1* 3' homology arm oligo 2 | *cgg*GTGCATTTGGCTGTCAGTATGGCCTCCAGACCGCCTCGACAGACA | Clone 3' 48 bp homology arm in GeneWeld vector for CRISPR-Cas9 targeted integration |
| 5'-junction-R | CGGTGGCTGAGACTTAATTACT | pPRISM genome/vector 5' PCR junction analysis [1] |
| 3'-junction-F | TTCAGATCAATTAACCCTCACC | pPRISM genome/vector 3' PCR junction analysis [1] |

1. Almeida, M.P., Welker, J.M., Siddiqui, S., Luiken, J., Ekker, S.C., Clark, K.J., Essner, J.J., and McGrail, M. (2021). Endogenous zebrafish proneural Cre drivers generated by CRISPR/Cas9 short homology directed targeted integration. Sci Rep *11*, 1732. 10.1038/s41598-021-81239-y.
