## Supplementary material for "Tracing developmental and adult hematopoiesis with an endogenous zebrafish *runx1-2A- CreERT2* CRISPR knock-in": Detailed Protocol

**Protocols for tamoxifen regulated CreERT2 tracing of Runx1 lineages in zebrafish *runx1-2A-CreERT2* embryos, larvae and adults.**

**Associated with manuscript:**

**Tracing developmental and adult hematopoiesis with an endogenous zebrafish *runx1-2A-CreERT2* CRISPR knock-in**

James A. Preston, Masuma K. Usha, Stephen C. Ekker, Karl J. Clark, Jeffrey J. Essner, Raquel Espin-Palazon, Maura McGrail

### A. Protocol for 4-OHT-induced lineage tracing in *runx1-2A-CreERT2* zebrafish embryos and larvae

**Summary:**

This protocol describes 4-hydroxytamixifen (4-OHT) treatment of zebrafish embryos or larvae for spatiotemporal induction of *runx1* lineage labeling with the zebrafish *runx1-2A-CreERT2* knock-in line. CRISPR knockin of *2A-creERT2* at the 3' end of the *runx1* coding sequence results in CreERT2 expression under the control of endogenous *runx1* gene regulatory elements. *runx1-2A-CreERT2* is combined with a floxed fluorescent reporter line to label the lineage of interest from the early embryo through larval stages. By controlling the timing of 4-OHT exposure during embryonic or larval development, recombination can be induced within defined hematopoietic waves and events, enabling analysis of Runx1 contributions to definitive hematopoiesis in the embryo and HSPC populations in larval hematopoietic organs. The protocol details genetic crosses, 4-OHT treatment regimen of embryos and larva, and sample preparation for live confocal imaging of fluorescently labeled Runx1 lineages during development.

Adhere to institutional protocols for laboratory safety and animal care and use.

**Timing:**

1-7 days depending on imaging stage

**Materials and Reagents:**

1. Zebrafish *Tg(runx1-2A-creERT2, exorh:CFPis97) (runx1-2a-creERT2)* (this paper)
2. Zebrafish *Tg(-3.5ubb:loxP-EGFP-loxP-mCherry)cz1701 (Mosimann et al., 2011) (ubi:Switch)*
3. Zebrafish *Tg(actb2:loxP-BFP-loxP-dsRed)sd27 (Bertrand et al., 2010)*

4. Nalgene petri dish (ThermoFisher 08-757-15C)
5. 28°C incubator
6. Aluminum foil
7. Nuclease-free 100% ethanol (EtOH)
8. E3 embryo media [1]
9. 0.06% 1-phenyl 2-thiourea (PTU) (Sigma P7629)
10. 0.4% Tricaine MS-222 Ethyl 3-aminobenzoate methanesulfonate (Sigma E10521)
11. 1.2% low melt agarose (Promega V2111)
12. Zeiss LSM 800 laser scanning confocal microscope or equivalent light microscope with Plan-Apochromat 10X/0.45 dry, 20X/0.8 dry, and W N-Achroplan 40x/0.75 water dipping objectives for fluorescence and differential interference microscopy/brightfield imaging
13. 10 mm 4-hydroxytamoxifen in 100% ethanol (4-OHT) (Sigma H6278)
14. Glass microscope slides (Fisher12550400) and No. 1 coverslips (Fisher 12541020)
15. FluoroDish coverslip bottom 35mm dish (World Precision Instruments FD350100) for live imaging with W N-Achroplan 40x/0.75 water dipping objective

**Procedure:**

1. Cross *runx1-2A-creERT2, exorh:CFPis97* with *ubi:Switch* or *actb2:loxP-BFP-loxP-dsRed* zebrafish.
2. Collect *runx1-2a-creERT2; ubi:Switch* or *runx1-2a-creERT2; ubi:Switch* embryos in 20 mL E3 embryo media in a petri dish and maintain at 28°C in an incubator.
3. At shield stage (6-8hpf), add 1 mL 0.06% PTU in E3 media to the dish for a final concentration of 0.003% to inhibit pigment synthesis.

<u>Continuous treatment with 4-OHT for CreERT2 induction from shield stage</u>:

4. Place up to 40 embryos in 20 ml E3 media in a petri dish.
   Add 10 μL 10 mm 4-OHT for a final concentration of 5 μM.
   Add 10 μL of 100% EtOH vehicle to control embryos.
5. Cover petri dish in aluminum foil and return to 28°C incubator.
6. Every 24 hours replace with fresh media containing 5 μM 4-OHT and 0.003% PTU and leave in induction media until desired stage for imaging.

<u>Short treatment with 4-OHT for timed induction of CreERT2 during development</u>:

7. Add desired timepoint place up to 40 embryos in 20 ml E3 media in pertri dish.
   Add 10 μL 10 mm 4-OHT for a final concentration of 5 μM.
   Add 10 μL of 100% EtOH vehicle to control embryos.
8. At end of timepoint induction window, discard treated E3 media and replace with fresh E3 media containing 0.003% PTU (foil covering is no longer required).

Imaging:

9. Place embryos or larva in petri dish in 30 ml E3 media containing 0.003% PTU. Anesthetize by adding 0.4%Tricaine to the media for a final concentration of 0.015%.
10. For imaging, mount zebrafish embryos or larvae in 1.2% low melt agarose/0.015% Tricaine on a slide or coverslip dish. Flood chamber with E3 media containing 0.003% PTU/ 0.015% Tricaine.

*runx1-2a-creERT2; ubi:Switch* embryos and larvae treated with 4-OHT during development can be raised to adulthood for long term lineage tracing of Runx1 hematopoietic precursors in adults. Raise *runx1-2a-creERT2; ubi:Switch* EtOH vehicle controls alongside 4-OHT treated embryos/larvae.

### B. Protocol for Adult 4-OHT-induced lineage tracing in *runx1-2A-CreERT2* zebrafish

**Summary:**

This protocol describes 4-hydroxytamixifen (4-OHT) treatment of adult *runx1-2A-creERT2; ubi:Switch* zebrafish for Runx1 lineage labeling in adult kidney marrow and peripheral blood. The protocol details genetic crosses, 4-OHT treatment regimen of adult fish, and collection of kidney marrow samples for downstream imaging and analysis.

Adhere to institutional protocols for laboratory safety and animal care and use.

**Timing:** 9 days

**Materials and Reagents:**

1. Adult *runx1-2A-creERT2; ubi:Switch* zebrafish
2. 0.8 L aquatic housing tank or equivalent container
3. 10 mm 4-hydroxytamoxifen in nuclease-free 100% ethanol (4-OHT) (Sigma H6278)
4. Nuclease-free 100% ethanol (EtOH)
5. 200 mL glass beaker
6. 0.4% Tricaine MS-222 Ethyl 3-aminobenzoate methanesulfonate (Sigma E10521)
7. Surgical equipment for removal of kidneys
8. Phosphate Buffered Saline (PBS) (0.8% NaCl, 0.02% KCl, 0.02 M PO4, pH 7.3)

**Procedure:**

1. Adult fish are maintained in tanks on a bench in fish facility for the duration of the experiment.
2. Place adult *runx1-2A-creERT2 ;ubi:Switch* zebrafish in 0.8 L tank filled with 200 mL system water.

3. Add 100 μL of 10 mm 4-OHT for a final concentration of 5 μM.
   Treat control fish by addition of 100 μL of 100% EtOH vehicle alone.
4. Protect treated tanks from light and incubate at 28°C for 12 hours overnight.
5. Remove treated water and replace with fresh system water for 12 hours.
6. Repeat steps 2-4 twice for a total of three cycles.
7. Remove water and replace with fresh system water, daily, for 6 days. Feed fish in the morning after water exchange. For afternoon feedings, food was added and 2 hours later fresh water exchanged.

Tissue collection for flow cytometry:

8. On day 9 place individual fish in 200 mL glass beaker with 100 ml system water.
9. Anesthetize fish by adding 0.4% Tricaine until final concentration exceeds 0.015%.
10. Place fish on surgical tray, collect peripheral blood as described [2,3], and excise kidney tissue.
11. Place extracted kidney tissue into ice cold PBS for dissociation and flow cytometry as described previously [2,3].

For long term Runx1 lineage tracing in adult kidney and blood, tissue collection can be performed on *runx1-2a-creERT2; ubi:Switch* adults treated as embryos or larvae with 4-OHT or vehicle control (Protocol A, above).
